# Predicting condensate formation of protein and RNA under various environmental conditions

**DOI:** 10.1101/2023.06.01.543215

**Authors:** Chin Ka Yin, Shoichi Ishida, Kei Terayama

## Abstract

**Motivation:** Liquid-liquid phase separation (LLPS) by biomolecules plays a central role in various biological phenomena and has garnered significant attention. The behavior of LLPS is strongly influenced by the characteristics of the RNAs and environmental factors such as pH and temperature, as well as the properties of the proteins. Recently, several databases of biomolecules associated with LLPS have been established, and prediction models of LLPS-related phenomena have been explored, leveraging these databases. However, a prediction model that concurrently considers proteins, RNAs, and experimental conditions has not been developed due to the limited information available from individual experiments in public databases.

**Results:** To address this challenge, we have built a new dataset called RNAPSEC, which serves each individual experiment as a data point. This dataset was accomplished by manually collecting data from public literature. Utilizing RNAPSEC, we developed two distinct models that consider a protein, RNA, and experimental conditions. The first model can predict the LLPS behavior of a protein and RNA under specific conditions. The second model can predict the required conditions for a given protein and RNA to undergo LLPS. RNAPSEC and these prediction models are expected to accelerate our understanding of the roles of proteins, RNAs, and environmental factors in LLPS.

**Availability:** The codes for the prediction models and RNAPSEC are available at https://github.com/ycu-iil/RNAPSEC.

## 1 Introduction

Liquid-liquid Phase Separation (LLPS) of biomolecules such as proteins and RNAs has attracted much attention due to their central role in various cellular phenomena and implications in several diseases. LLPS is a physicochemical process that allows the formation and maintenance of condensates composed of specific biomolecules (Hyman, Weber and Jülicher 2014; Boeynaems *et al*. 2018). These condensates exhibit liquid-like properties and can respond to specific extracellular and intracellular signals by fusing, exchanging surrounding components, or dissolving (Youn *et al*. 2019). In addition to the properties of proteins, the formation and maintenance of LLPS are regulated by RNAs and multiple environmental factors, such as, surrounding pH, and temperature (Li *et al*. 2022). Dysregulation of LLPS has been suggested to induce a phase transition from liquid-like condensates to solid-like condensates, leading to the formation of aggregates and amyloids (Shin and Brangwynne 2017; Wang *et al*. 2021). These aggregates and amyloids are characteristic features found in neurodegenerative diseases such as Alzheimer’s disease and Amyotrophic Lateral Sclerosis (ALS) (Murakami *et al*. 2015; Ambadipudi *et al*. 2017). Given that the LLPS behavior is strongly associated with cellular phenomena and several diseases, it is important to evaluate the LLPS behavior of biomolecules to elucidate their function and disease relevance.

In recent years, there has been an increase in publicly available databases, and research utilizing machine learning models has intensified (Li *et al*. 2020; Mészáros *et al*. 2020; Ning *et al*. 2020; Raimondi *et al*. 2021; van Mierlo *et al*. 2021; Chen *et al*. 2022; Chu *et al*. 2022; Liu *et al*. 2022; Wang *et al*. 2022; Zhu *et al*. 2022). Previous models have been employed to identify related proteins and demonstrate the effectiveness of these approaches. These models use only the sequence or sequence-derived properties of a protein as input and output the LLPS behavior (van Mierlo *et al*. 2021; Chen *et al*. 2022; Chu *et al*. 2022). Furthermore, a prediction model has been developed that takes a protein sequence and experimental conditions as input and outputs the propensity of the protein to undergo LLPS, using the LLPS-related database LLPSDB (Li *et al*. 2020; Raimondi *et al*. 2021). Through dataset analyses and assessments of feature importance in prediction models, these studies have identified protein features that significantly influence LLPS behavior. Therefore, the machinelearning based approach holds potential for enhancing our comprehensive understanding of LLPS behavior.

However, a prediction model that considers proteins, RNAs, and experimental conditions has not been developed, despite their recognized significance in the regulation of LLPS (Alberti, Gladfelter and Mittag 2019). Recent studies have shown additional insights into the roles of RNAs and environmental factors in LLPS regulation (Roden and Gladfelter 2021). Numerous RNAs have been discovered within membraneless organelles, and it is suggested that various properties of RNAs, including their concentration, length, structure, and sequence, can influence the behavior of LLPS (Garcia-Jove Navarro *et al*. 2019; Grese *et al*. 2021; Henninger *et al*. 2021; MATSUI and NOZAWA 2021; Roden and Gladfelter 2021; Wiedner and Giudice 2021). Additionally, *in-vitro* experiments have reported that changes in environmental factors, such as pH, temperature, and ionic strength, can also alter the behavior of LLPS (Ambadipudi *et al*. 2017; Rayman, Karl and Kandel 2018; Gui *et al*. 2019; Li *et al*. 2022). Therefore, to better understand LLPS as a biological phenomenon, it is desirable to consider proteins, RNAs, and experimental conditions.

In this study, we aimed to develop a prediction model that considers proteins, RNAs, and experimental conditions. Since the performance of a machine-learning model is heavily influenced by the quality and quantity of a training dataset, having more detailed information is desirable. However, the experimental information corresponding to a single experiment is not available in public databases due to inconsistent recording formats, missing values, and the use of range and multiple notations to record information from multiple experiments as a single data point. Therefore, we first constructed a dataset, RNAPSEC (RNAPhaSep with detailed Experimental Conditions), with single experiments as entries by thoroughly reviewing the public literature resisted in RNAPhaSep (Zhu *et al*. 2022), which contains data about LLPS-related RNAs, and manually collecting detailed experimental information from experiments conducted by a protein and RNA. Using RNAPSEC, we successfully developed a model to predict whether a given protein and RNA can undergo LLPS under specific conditions. This model has shown a relatively high performance with a ROC-AUC of 0.69 in cross-validation. By using this model, we were able to construct phase diagrams that reflect experimental results by predicting LLPS behaviors under various experimental conditions. In addition, we also developed a model that predicted the experimental conditions required for a given protein and RNA to undergo LLPS. This model also showed a relatively high performance with a ROC-AUC exceeding 0.60 for each experimental condition. These results suggest that both models are effective in predicting LLPS behavior and conditions, indicating potential applications in understanding the mechanisms of LLPS.

## 2 Methods

The overview of this study is illustrated in Figure 1. The first step was to construct a dataset, RNAPSEC, which involved manually collecting experimental conditions from the public literature resisted in RNAPhaSep. The second step was to construct the prediction models, which can be divided into two phases: data processing and model development. In the data processing phase, we selected the needed data for the model development from RNAPSEC and processed them for feature extraction from the experimental conditions, the protein sequence, and the RNA sequence. In model development, we created a model that takes protein sequence-derived features, RNA sequence-derived features, and experimental condition-derived features as inputs and outputs whether the LLPS will occur or not. Additionally, we developed a model that takes protein sequence-derived features and RNA sequence-derived features as inputs and outputs the experimental conditions required for a protein and RNA to undergo LLPS.

**Figure 1.**
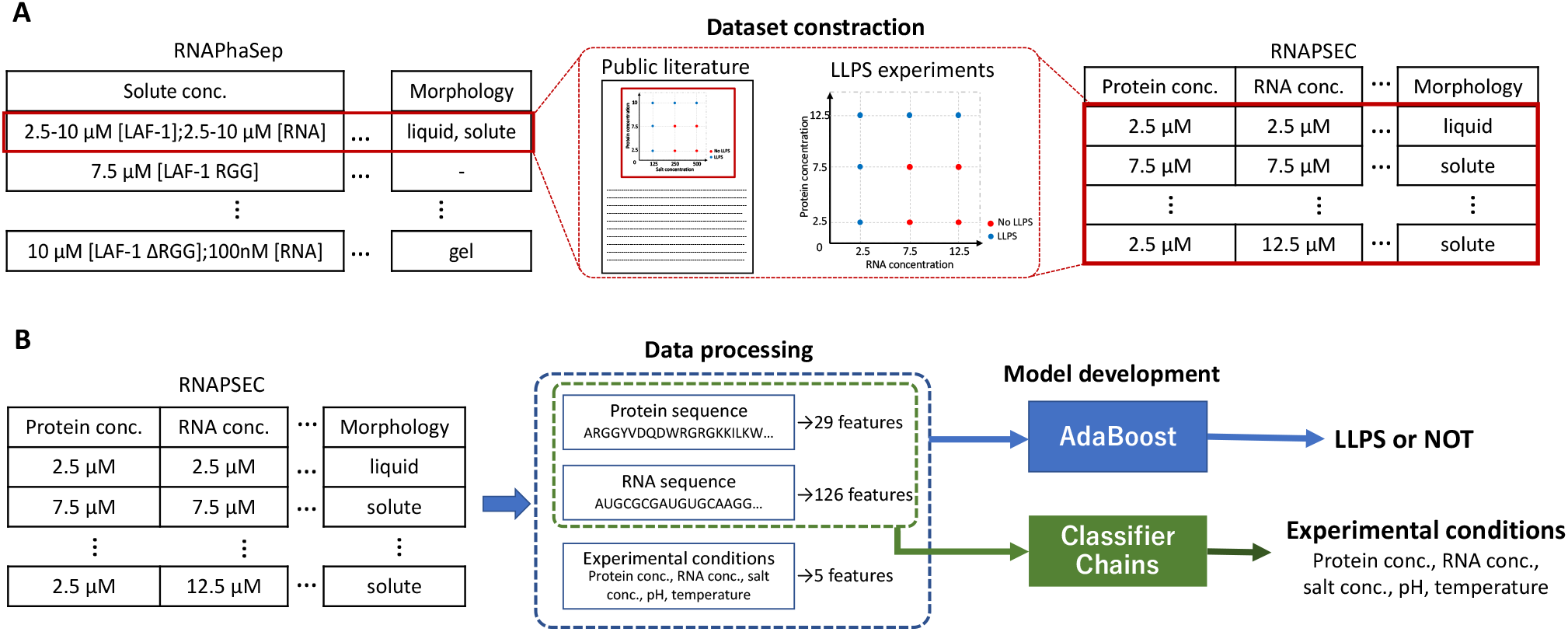
Overview of the construction process for RNAPSEC dataset and the machine-learning-based prediction models. We first constructed the dataset RNAPSEC (A) and subsequently performed the model construction (B). (A) Overview of the dataset construction. The experimental information from RNAPhaSep, where multiple experiments were grouped together, was disassembled into individual experiments. To achieve this, we manually extracted the experimental information from the public literature stored in RNAPhaSep and recorded one experiment as one entry. (B) An overview of the model construction. We constructed two distinct models trained using RNAPSEC. The first model predicts LLPS behavior using 29 protein-derived features, 126 RNA-derived features, and 5 experimental conditions. The second model predicts the required conditions for a protein and RNA to undergo LLPS using 29 protein-derived features and 126 RNA-derived features.

### 2.1 Construction of RNAPSEC

RNAPSEC was constructed by gathering experimental information from relevant papers (Figure 1A). To collect data, we selected experiments that utilized a single protein and RNA from the public literature referenced in RNAPhaSep. We manually extracted the experimental information, including protein concentration, RNA concentration, salt concentration, buffer pH, temperature, and experimental results. To simplify the data processing, protein concentration, RNA concentration, and salt concentration were recorded in a tabular format with separate values and units, while only values for temperature and pH were recorded. Additional experimental parameters, such as incubation time and small molecules, along with information about biomolecules, were recorded identically to RNAPhaSep. Experimental results were recorded as described in the respective articles. In the absence of descriptions, results were inferred from the size and shape of microscopic images. Microscopic images that showed no distinctive features were classified as solutes, images with only spherical granules were classified as liquid-like condensates, images with reticulated networks were classified as gel-like condensates, and images with isolated objects that did not take on a spherical shape were classified as solid-like condensates. Data for which the result could not be determined from either the text or the images were excluded.

### 2.2 Preprocessing process of experimental conditions and feature extraction from sequences

For the data analysis and the model development, we standardized the units and descriptions across all data. Data with the following features were selected for processing: usage of a single protein and RNA, without any deficiency in sequence information, experimental results, or experimental conditions. Protein concentration, RNA concentration, salt concentration, pH, and temperature were extracted from RNAPSEC. Protein concentration and RNA concentration were unified as μM and converted into common logarithms. Salt concentration was converted to ionic strength using the pyEQL package (R.S. Kingsbury 2013). Data using the salt not supported by pyEQL were excluded. The temperature was unified to °C, with room temperature (RT) represented as 25 °C. For pH, values were used directly. After completing all processing steps, data with missing values were excluded. To compare the distributions of experimental conditions between RNAPSEC and RNAPhaSep, protein concentration and RNA concentration in RNAPhaSep were preprocessed in the same way. However, as the protein concentration and RNA concentration were recorded in a single column in RNAPhaSep, data with the following descriptions were excluded: data where it was unclear whether the described concentration referred to a protein or an RNA, where either concentration was not recorded, or where multiple experimental results were described. If a range notation or multiple values were mentioned, the average was used. Protein sequences were converted into single-letter amino acid sequences and then transformed into features showing various properties such as amino acid composition, hydrophobicity, and isoelectric point using Biopython (Cock *et al*. 2009). Detailed descriptions of the features are provided in Table S1. Similarly, RNA sequences were converted into single-letter nucleotide sequences and transformed into features such as nucleotide composition and sequence periodicity using MathFeature (Bonidia *et al*. 2022). Detailed descriptions of the features are provided in Table S2. A total of 131 features were used, including 5 from experimental conditions, 29 from a protein sequence, and 97 from an RNA sequence, as input features for the model.

### 2.3 Construction of prediction models and their validations

In this study, we trained and evaluated two models with distinct inputoutput features. The first model predicts whether a protein and RNA will induce LLPS under specified experimental conditions, based on an input of 29 features from a protein sequence, 97 features from an RNA sequence, and 5 features from experimental conditions (Figure 1B). We trained and evaluated using several models with different algorithms: a Logistic Regression (LR), a K-Nearest Neighbor (KNN), a Gaussian Naïve Bayes (GaussianNB), a Random Forest (RF) (Ho 1995), a Light Gradient Boosting Machine (LightGBM) (Ke *et al*. 2017) and an Adaptive Boosting (AdaBoost) (Schapire 2013). All models except the LightGBM model were conducted using the scikit-learn library in Python (Pedregosa *et al*. 2011). The models, except for the decision tree-based models, were trained and evaluated using the data standardized to have a mean of 0 and a variance of 1. The performances of these models were assessed by splitting data into a training dataset and test dataset using stratified group 10fold cross-validation. The groups for the stratified group 10-fold crossvalidation were assigned according to the protein sequence of each data. Within each fold, hyperparameters of the models were adjusted to maximize the ROC-AUC of the test dataset (Table S3). The average of a ROCAUC and feature importance was calculated from each fold, and the final model was trained on the full dataset.

Furthermore, phase diagrams were constructed using the model trained in each fold of the cross-validation. For each fold, we prepared a new test dataset with extended protein concentration and RNA concentration for each protein-RNA pair. The range of protein concentration and RNA concentration in the test dataset was expanded by an interval of 0.2, 1 less than the minimum and 1 greater than the maximum. We made the predictions on the expanded test dataset using the trained model and plotted phase diagrams for each protein-RNA pair.

The second model predicts the experimental conditions for a protein and RNA to undergo LLPS, based on the inputs of 29 features from a protein sequence, and 97 features from an RNA sequence (Figure 1B). Since the LLPS behavior is not determined by a single environmental factor, we used the classifier chains to develop the model. The Classifier Chains are multiple classifiers connected in series, which allows classification to be performed by including the prediction results of one previous classifier as input (Read *et al*. 2011). Therefore, predictions can be carried out by considering multiple experimental conditions. We trained the classifier chains based on the AdaBoost model, which showed the best performance in the model predicted the LLPS behavior, with the hyperparameters below: decision tree classifier without setting max depth for the base estimator, learning rate for 0.5, random state for 116. Each experimental condition was classified into several classes according to its value and treated as a classification problem. pH was classified into three classes: acidic, neutral, and basic; temperature into three classes: low temperature, room temperature, and high temperature; protein concentration, RNA concentration, and ionic strength into five classes according to 20 %, 40 %, 60 %, 80 % and 100% of the value distribution (Table 1). A ROC-AUC and confusion matrices were calculated from the predicted results of group 10-fold crossvalidation. The groups of the cross-validation were assigned by the protein sequence of each data. The final model was trained by a full dataset.

**Table 1.** Classification of experimental values into classes. Each experimental condition was classified into three or five classes depending on its value.

| Class | 1 | 2 | 3 | 4 | 5 |
| --- | --- | --- | --- | --- | --- |
| pH | 0-7 | 7-8 | 8-14 | - | - |
| Temperature | 0-25 | 25-28 | 38- | - | - |
| Ionic strength | 0.00-0.03 | 0.03-0.07 | 0.07-0.15 | 0.15-0.18 | 0.18- |
| Protein conc. | -4.60--0.75 | -0.75--0.90 | -0.90-2.03 | 2.03-3.71 | 3.71-6.22 |
| RNA conc. | -10.8--8.03 | -8.03--4.93 | -4.93--1.84 | -1.84-0.87 | 0.87-6.62 |

## 3 Results

### 3.1 Data content in RNAPSEC

RNAPSEC contains a total of 1563 entries, including 394 solute data without LLPS, 1023 liquid data with liquid-like condensates, 92 gel data with gel-like condensates, and 54 solid data with solid-like condensates. These data points originate from 30 proteins with 82 unique sequences and 134 RNAs. In RNAPhaSep, experiments with a single protein and RNA, including data without or more than one description of experimental results (described as “Unknown” and “Others” in Figure 2A). RNAPSEC, meticulously curated with data from public literature, has been designed to avoid the inclusion of ambiguous entries where experiments involving a single protein and RNA might lack the experimental result with clear descriptions or might have more than one description. In terms of protein concentration and RNA concentration, RNAPSEC exhibits a broader distribution compared to the more scattered one seen in RNAPhaSep (Figure 2B, C). Consequently, RNAPSEC is equipped to provide a more comprehensive understanding of the conditions and results of each experiment.

**Figure 2.**
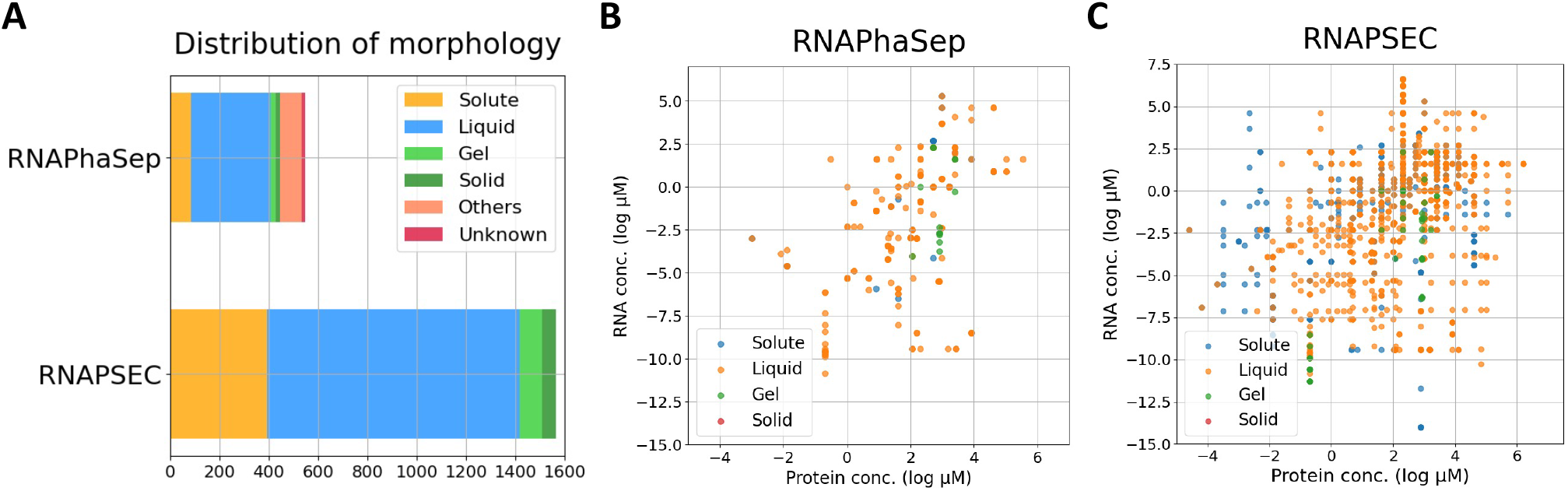
Distribution of phase behavior and protein-RNA concentrations in RNAPShaSep and RNAPSEC. (A) Distribution of phase behaviors outcomes from each experiment. (B) Distribution of protein and RNA concentrations in preprocessed RNAPhaSep. (C) Distribution of protein and RNA concentrations in preprocessed RNAPSEC.

Previous studies have shown that changes in protein concentration and RNA concentration can alter the LLPS behavior (Li *et al*. 2022), suggesting a possible trend between concentration changes and LLPS behavior. However, no obvious tendencies were identified from the distribution of protein concentration and RNA concentration in RNAPSEC (Figure 2C). These results suggest that the LLPS is a complex phenomenon determined by multiple experimental parameters and properties of biomolecules. It is also likely that the lack of trends from the distribution is due to the limited number of data points.

### 3.2 Evaluation of the model that predicts the LLPS behavior of a protein and RNA under given conditions

We trained and evaluated the model to predict the LLPS behavior using 1381 data recorded in RNAPSEC, including 387 solute data as negative data and 994 liquid data as positive data. This model takes 126 features of the sequence-derived features and 5 features of the experimental conditions as inputs and outputs on whether a given protein and RNA can undergo LLPS under certain conditions. We compared the performances of the prediction models using different five algorithms, LR, KNN, GaussianNB, RF, LightGBM, and AdaBoost, and selected the best. The performance of each model was estimated by stratified group 10-fold cross-validation with protein sequences as group labels. As a result, the decision tree-based models (the LightGBM model, the RF model, and the AdaBoost model) showed superior performances, with the AdaBoost model exhibiting the highest ROC-AUC of 0.69 (Figure 3A, Figure S1). In addition, the AdaBoost model had a precision of 0.78, indicating its ability to accurately classify approximately 78% of LLPS experiments with liquidlike condensates. Hence, this model can effectively predict the LLPS behavior of a protein and RNA under specific conditions and can be used to screen the required conditions to undergo LLPS. We also calculated the feature importance in the AdaBoost model and found that protein concentration and RNA concentration were significantly more important than the other features (Figure 3B). It suggested that the LLPS behavior may be heavily influenced by these features.

**Figure 3.**
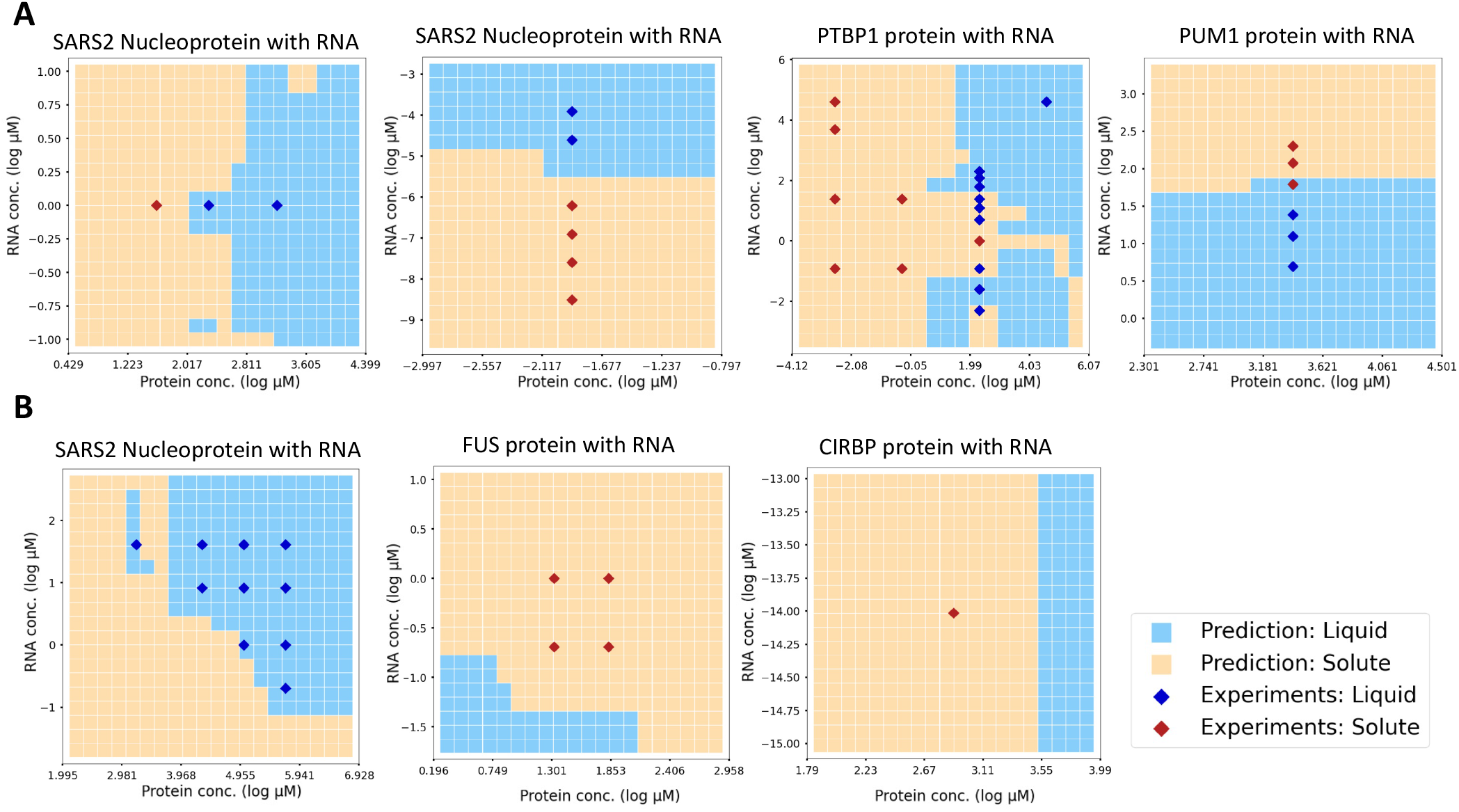
Examples of phase diagram depictions using the model predicting LLPS behavior. Phase diagrams were constructed from the predicted results of the expanded test dataset with varying protein concentration and RNA concentration at regular intervals. (A) Examples of phase diagrams reflecting experimental results. (B) Examples of new behavior changes were predicted to occur with shifting input concentration.

Although our model showed a relatively high ROC-AUC of 0.69, it was slightly lower than the previous models that only considered a protein sequence (van Mierlo *et al*. 2021; Chu *et al*. 2022). This result might be due to the model considering three different factors: a protein, RNA, and experimental conditions, which renders the prediction problem more complex than considering a protein alone. However, given the complexity of LLPS behavior — a phenomenon influenced by the interplay of at least these three factors — it’s necessary for prediction models to account for all these parameters. To improve the performance of our model and allow for more detailed analyses, we need to expand RNAPSEC with unique sequences and conditions in future studies.

### 3.3 Construction of phase diagram

The prediction model developed in this study can immediately predict the LLPS behavior of a single protein and RNA, and is therefore useful for the rapid construction of a phase diagram showing the LLPS behavior under various conditions. To evaluate the validity of these phase diagrams constructed using the prediction model, we extended the range of protein concentration and RNA concentration used in the experiments from a minimum of minus 1 to a maximum of plus 1, with increments of 0.2, while keeping other experimental conditions constant.

Figure 4 shows the phase diagrams constructed by the trained model. In each figure, the red and blue squares represent experimental results, with red indicating that LLPS did not occur and blue indicating that liquid-like condensates were observed. The orange and light blue circles represent the prediction results; orange represents the absence of LLPS, and light blue represents the formation of liquid-like condensates. As a result, the orange circles were more common in areas with the red squares and the light blue

**Figure 4.**
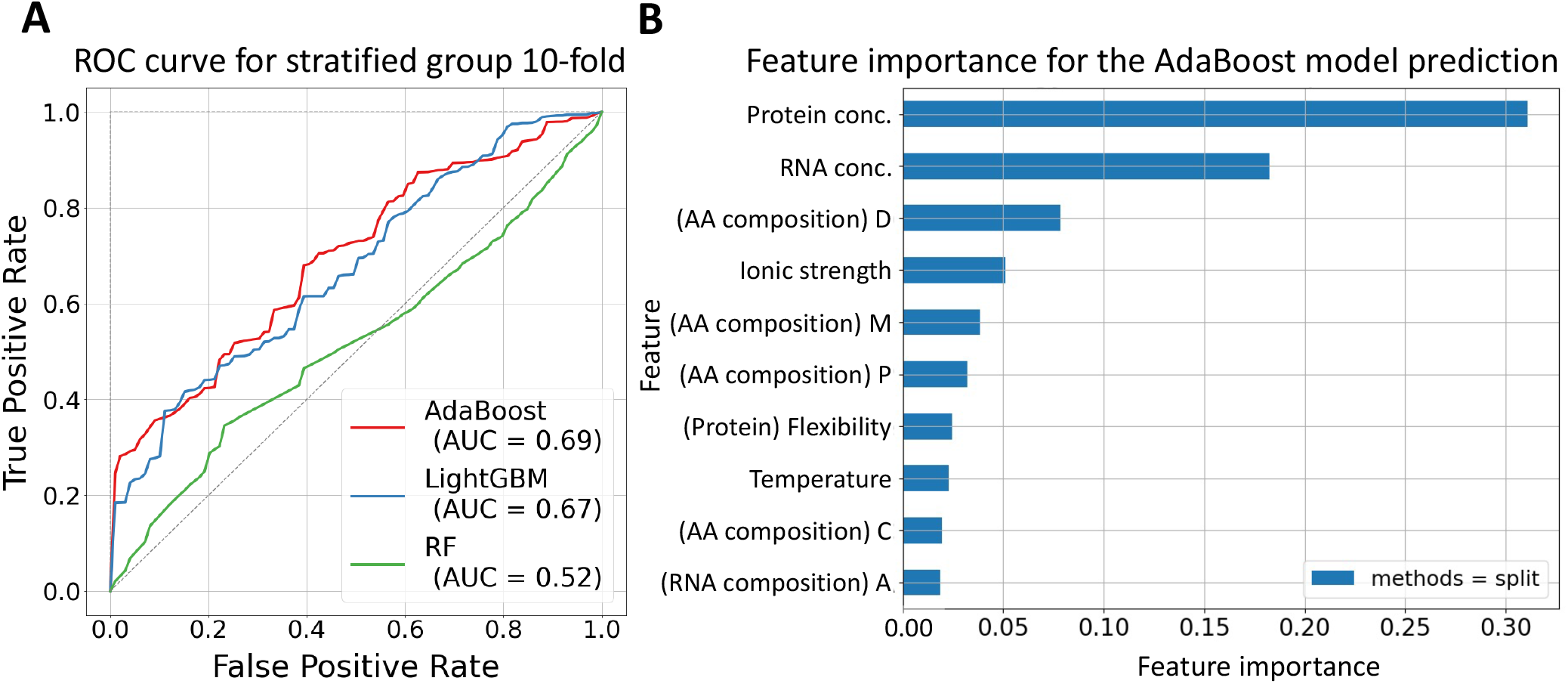
Performance and feature importance of machine-learning-based models in predicting LLPS behavior. (A) The ROC curve of the RF model, the LightGBM model, and the AdaBoost model with the stratified group 10-fold cross-validation. Each line represents the average line of the ROC curve for each fold in cross-validation. (B) The top 10 effective features for the prediction through the AdaBoost model were shown by calculating their feature importance.

### 3.4 Development of the model that predict required conditions for a protein and RNA to undergo LLPS

Furthermore, we developed a model capable of predicting the required conditions for LLPS using the classifier chains based on the AdaBoost models. The model was trained and evaluated using 994 liquid data from RNAPSEC. The model takes the features derived from a protein sequence and RNA sequence as inputs and outputs experimental conditions including RNA concentration, protein concentration, ionic strength, pH, and temperature. The performance of this model was evaluated using Group 10-Fold CV and scored with ROC-AUC and confusion matrices. As a result, the ROC-AUC was higher than 0.60 for all five experimental conditions (Figure 5A). However, it was difficult to accurately predict experimental classes in protein concentration, RNA concentration, and ionic strength (Figure 5B).

**Figure 5.**
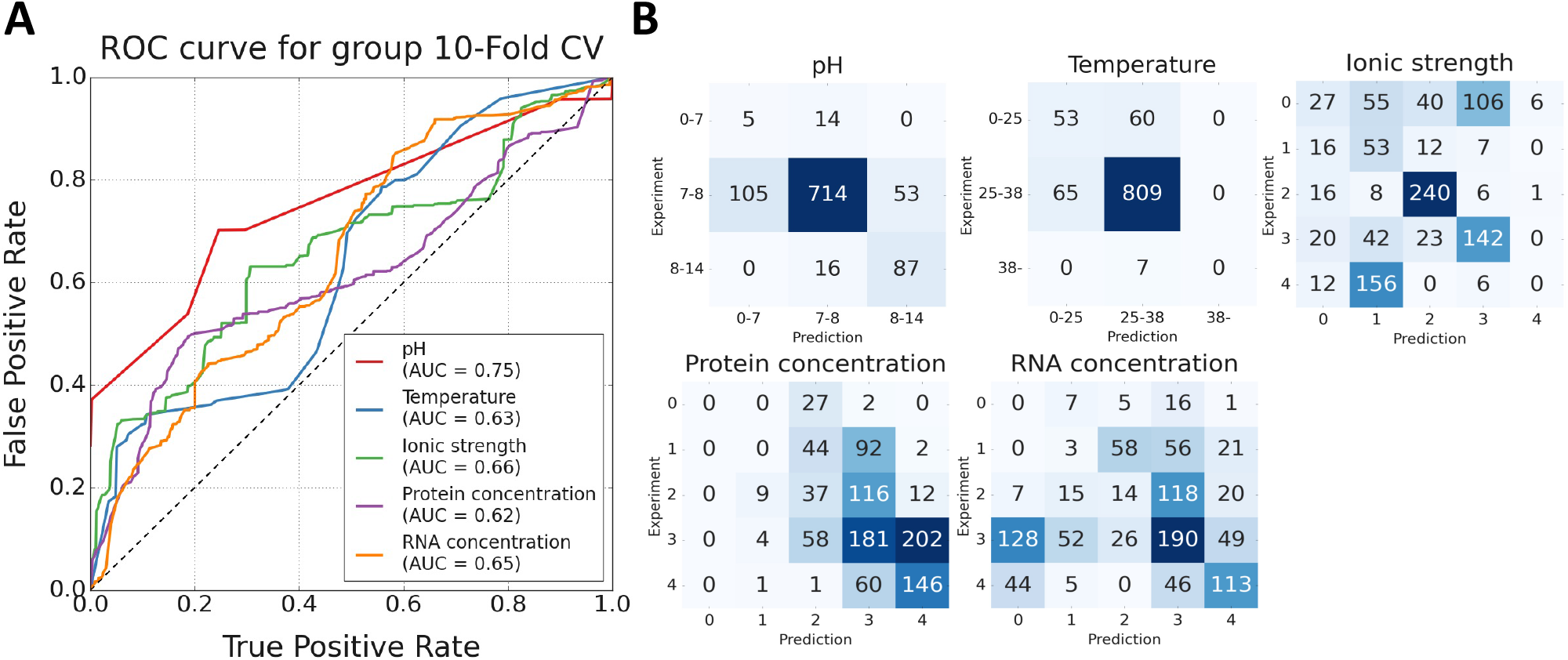
Performance of classifier chains model with AdaBoost models in predicting five experimental conditions. (A) The ROC curve of the model predicts the required conditions with group 10-fold cross-validation. Each line represents the average ROC curve of each fold in the crossvalidation. (B) Confusion matrices for the models predicting each experimental condition with the group 10-fold cross-validation. circles in areas with the blue squares. This suggested that the predicted results aligned well with the experimental outcomes (Figure 4A). Intriguingly, in experiments where only solutes were observed, the LLPS behaviors were predicted to be altered by shifting the input value of protein concentration and RNA concentration (Figure 4B). This suggested that the LLPS behavior may have the potential to change as predicted results under given conditions. In summary, using this model can overlook the LLPS behaviors under various conditions and enable a comprehensive screening for the required conditions to undergo LLPS.

## Conclusion

In this study, we have developed a model that predicts the LLPS behavior, considering both RNA and experimental conditions in addition to a protein. For this purpose, we constructed a dataset, RNAPSEC, where each experiment forms a data point. RNAPSEC contains a total of 1546 data with a large amount of information from various LLPS experiments on proteins and RNAs. In addition, we developed the model to predict the LLPS behavior of a protein and RNA under specific conditions. The phase diagrams constructed by this model reflected the experimental results and could be effective for large-scale screening of LLPS conditions. However, the model that predicts the experimental conditions for a protein and RNA to undergo LLPS required further improvements in the model performance for practical application. The application of these tools may open new avenues for future research, enabling the development of more accurate predictive models and broadening our perspective on the complex interplay between proteins, RNAs, and environmental factors in the LLPS by biomolecules.

