## Supplemental Information for "Predicting condensate formation of protein and RNA under various environmental conditions"

* To whom correspondence should be addressed.

### S1 Supplementary Tables

**Table S1. Feature extraction from a protein sequence.** The features described in Table S1 were extracted from a protein sequence using Biopython and used as inputs for the models described in the text. The descriptions below were referred in (Cock *et al.* 2009).

| Feature | Detail | Dimension |
|---|---|---|
| Amino acids composition | The composition of each amino acid. | 20 |
| Molecular weight | The molecular weight of a protein | 1 |
| Gravy | Calculated the hydrophobic of a protein (Kyte and Doolittle 1982). | 1 |
| Aromaticity | The relative frequency of F, W, Y. | 1 |
| Instability | Calculated the stability of protein using the method of Guruprasad *et al.* (Guruprasad, Reddy and Pandit 1990). | 1 |
| Flexibility | Calculated the average flexibility of a full-length protein using the method of Vihinen *et al*. (Vihinen, Torkkila and Riikonen 1994). | 1 |
| Isoelectric point | Return the isoelectric point of a protein | 1 |
| Secondary structure fraction | Return the fraction of amino acids below.<br>- Amino acids in helix: V, I, Y, F, W, L.<br>- Amino acids in turn: N, P, G, S.<br>- Amino acids in sheet: E, M, A, L. | 3 |

**Table S2. Features extraction from an RNA sequence.** The features described in the Table S2 were extracted from an RNA sequence using MathFearture and used as inputs for the models described in the text. The descriptions below were referred in (Bonidia *et al.* 2022).

| Feature groups | Feature | Dimension |
|---|---|---|
| Nucleic acid composition | Nucleic Acid composition | 4 |
| | Dinucleotide composition | 16 |
| | Trinucleotide composition | 64 |
| Entropy | Shannon entropy | 2 |
| | Tsaillis entropy | 2 |
| Fourier transform | Z-curve + Fourier | 2 |
| | Real + Fourier | 2 |
| | Binary + Fourier | 2 |
| Open reading frame (ORF) | The average length of ORF | 1 |
| Fickett score | Fickett score calculated from a full-length and ORF. | 2 |

**Table S3. Hyper-parameter tuning for the models to predict the LLPS behavior.** Table showing the parameters configuration and tuning ranges for hyper-parameter tuning. We use the grid search to tune the models except LightGBM model. The hyper parameters of the LightGBM model have been tuned with 50 trials using Optuna (Akiba *et al.* 2019). The hyper-parameters of the other models were tuned using grid search.

| Model | Parameter | Value/Value Range |
|---|---|---|
| LightGBM | lambda l1 | $1.00\times10^{-8}$ - 10 |
| | lambda l2 | $1.00\times10^{-8}$ - 10 |
| | Max number of leaves | 2 - 256 |
| | Feature fraction | 0.4 - 1 |
| | Bagging fraction | 1 - 7 |
| | Min child samples | 5 - 100 |
| RF | Number of estimators | 10, 100, 200, 300, 400, 500 |
| | Criterion | Gini, entropy |
| | Max depth | 1, 2, 3, 4, 5 |
| | Max feature | sqrt, log2, None |
| AdaBoost | Number of estimators in the Decision tree as base estimator. | 5, 6, 7, 8, 9, 10 |
| | Learning rate | 0.5, 1, 1.5 |
| GaussianNB | Variance smoothing | Log (-9) - 1 |
| LR | C | $1.00\times10^{-5}$ - $1.00\times10^{5}$ |
| | Random state | 0-101 |
| KNN | Number of neighbors | 1- 21 |
| | Weights | Uniform, distance |
| | Power parameter for the Minkowski metric | 1, 2 |

S2 Supplementary Figure

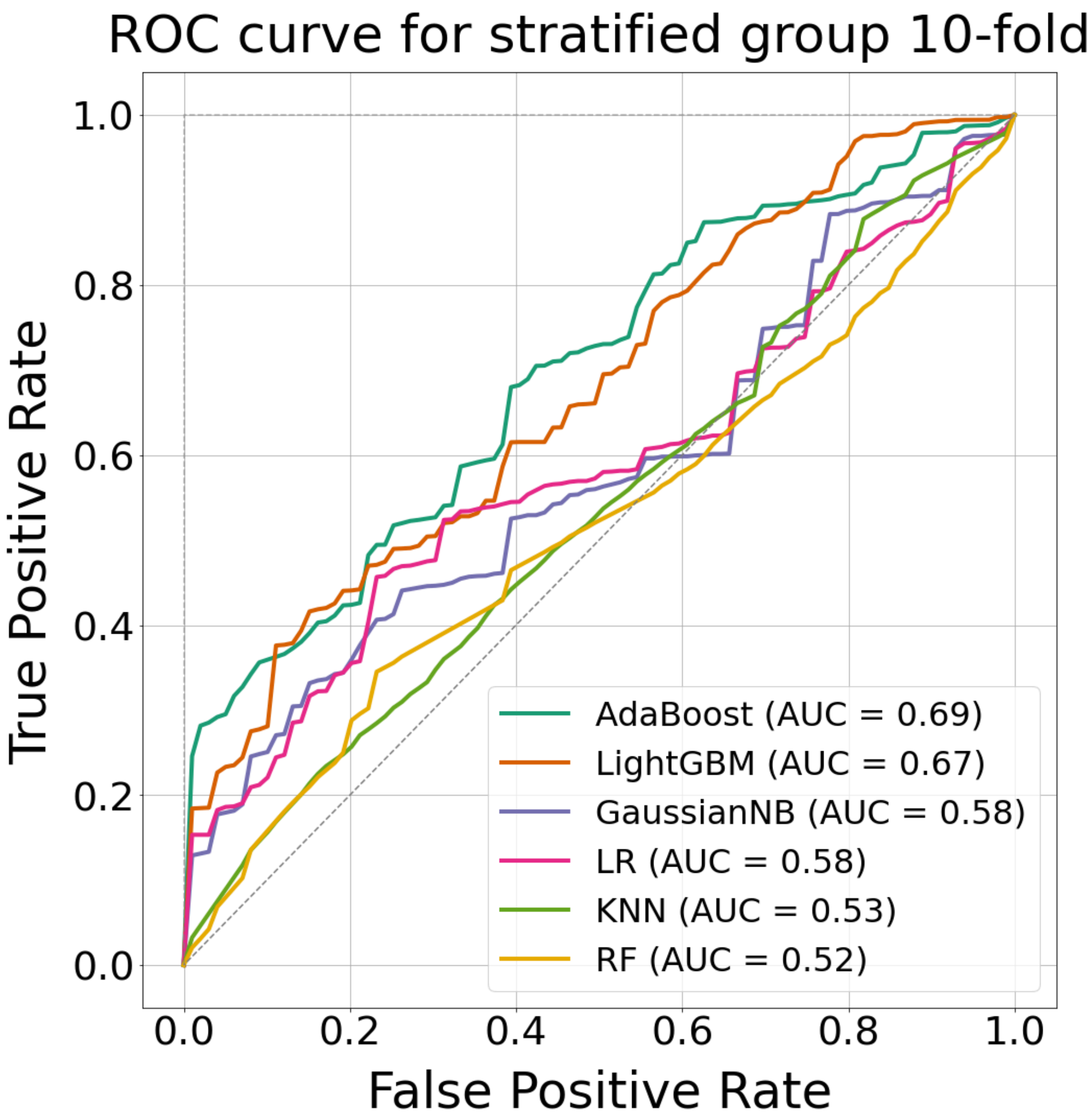


**Figure S1. The ROC curve for the stratified group 10-fold cross-validation of the models predicting LLPS behavior**. Figure showing the ROC curve obtained with the stratified group 10-folds cross-validation of the models that predict the LLPS behavior of a given protein and RNA described in the paper.
